# Ultrasensitive Regulation in the Tryptophan System in *Escherichia coli* Provides Effective Noise Mitigation

**DOI:** 10.64898/2026.11.02.763317

**Authors:** Vinay A. Bavdekar, Sharad Bhartiya, K. V. Venkatesh

## Abstract

The tryptophan system in *Escherichia coli* maintains intracellular concentration of tryptophan in a narrow band regardless of the nutritional status and uncertainty in its environment. The system is characterised by three distinct regulations namely, genetic regulation, mRNA attenuation and enzyme inhibition, that use the tryptophan concentration as a feedback signal. It has been shown previously, that the multiple feedback loop design is responsible for rapid synthesis of tryptophan when faced with starvation. It is well known that a design with rapid response to external signals also implies poorer noise mitigation properties. This work investigates the design elements that limit the adverse impact of the multiple feedback loop architecture on noise propagation in the tryptophan system using a frequency response analysis. Specifically, we use sinusoidal input describing functions to approximate the transfer characteristics of the ultrasensitive regulators and then obtain the frequency response of the closed-loop feedback system. The study suggests that the tryptophan system of *E. coli* has evolved to simultaneously satisfy the two properties of rapid response and noise mitigation. Particularly, the multiple feedback loop is responsible for the rapid response, while the nonlinear regulation provides effective noise mitigation.

## 1 Introduction

Living systems operate efficiently and in a stable manner to meet various survival-related objectives in an uncertain environment. Such systems must be robust and must provide an appropriate physiological response in the presence of these uncertainties[2,11]. To this end, nature has evolved structural motifs including cascades[19], feedforward control[4,15,20,26] and multiple feedback loops[14] resulting in complex regulatory networks[1,8,12,13,23]. A systems level analysis of these complex networks is complicated due to numerous interactions, which in turn results in a large number of system parameters. Ultimately, the phenotypic response of the network is a manifestation of the design principles resulting from the structural motif and system parameters[9]. It is, therefore, of interest to elucidate the design principles and relate them to the phenotypic response.

The tryptophan system of *Escherichia coli* has been well characterized and therefore amenable to a systems level analysis[6,24]. The primary objective of the tryptophan system in *E. coli* consists of maintaining the concentration of tryptophan, an essential amino acid, in a narrow band regardless of its nutritional status. Thus, in the absence of tryptophan in the medium, *E.coli* exhibits a rapid synthesis of tryptophan to ensure its survival. On the other hand, in a tryptophan rich medium, the synthesis is shut off and the uptake is regulated to maintain the desired concentration, thereby avoiding toxicity on account of excess intracellular concentration. Moreover, the tryptophan system must achieve its goals in an uncertain environment (extrinsic noise) as well as in presence of randomness in molecular interactions (intrinsic noise)[16,22]. In this work, we analyse the role of the regulator in providing robustness to noise while responding rapidly to nutritional demands.

## 2 The tryptophan system

Tryptophan is synthesized from chorismate, an intracellular nitrogen source with the aid of four distinct catalysing enzymes[7,17,18] (see Fig. 1). These four enzymes namely anthranilate synthase (AS), phophoribosyl anthranilate transferase (PRT), indole glycerol phosphate synthase (IGP) and tryptophan synthase (TS) are encoded from five distinct structural genes namely *trpE, trpD, trpC, trpB* and *trpA* of the tryptophan operon. However, the synthesis of the enzyme depends on the state of the operon, repressed or induced. As seen from Fig. 1, the operator overlaps the promoter and hence repressor and RNA polymerase molecules compete to determine the initiation of transcription. Two molecules of intracellular tryptophan (*T*) bind to the aporepressor (*A*_*p*_) that is transcribed from the gene *trpR* adjacent to promoter/operator site. This active complex of tryptophan-aporepressor called the holorepressor (*Hp*), subsequently binds to the operator site. Once the operator is bound to the holorepressor, RNA Polymerase (RNAP) can no longer bind to the promoter site, thus preventing initiation of transcription. This is the first stage in regulation of the tryptophan synthesis and this mechanism is termed as genetic repression *C*_1_(*T*). The second stage in regulation of the tryptophan synthesis is transcriptional attenuation *C*_2_(*T*). The attenuator is a leading region, which is located between the promoter and the structural genes. This region is made up of four equally large segments, which may fold to form three different RNA secondary structures or hairpins. The hairpin formed from the first two transcribed segments may stall the polymerase from further transcription. A ribosome binds to the hairpin structure to initiate translation, which disrupts the hairpin thereby enabling further transcription. At low tryptophan concentrations, a hairpin comprising of the second and third segments is formed, which is recognized as an anti-terminator resulting in transcription of structural genes. On the other hand, at high tryptophan concentrations, the third and fourth segments of the attenuator region form the hairpin structure, which is recognized as a terminator thus destabilizing the polymerase-DNA complex and prematurely aborting the transcriptional process. Attenuation thus mainly contributes in the termination of the transcription process that leads to incomplete mRNA, which cannot synthesize the structural enzymes. In the third level of regulation, excess tryptophan molecules bind to the enzymes synthesized by structural genes and turn off their activity. This mechanism is termed as enzyme inhibition *C*_3_(*T*). This is shown by the dashed oval encompassing all the catalysing enzymes of the pathway in Fig. 1.

**Fig. 1.**
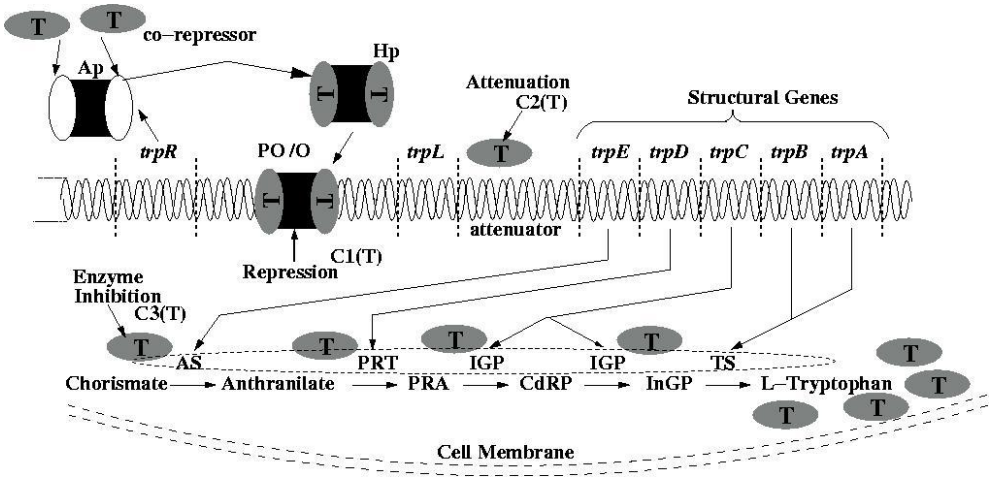
Schematic of the tryptophan synthesis pathway

The molecular description of the tryptophan system discussed above can be abstracted as three processes in series namely transcription, translation, and synthesis[5,24] (see Fig. 2). The regulatory structure of the tryptophan system senses the intracellular tryptophan concentration and regulates at three distinct levels[25] through genetic regulation, transcriptional attenuation and enzyme inhibition thereby constituting a multiple feedback loop structure (*C*_1_, *C*_2_ and *C*_3_ in Fig. 2, respectively). Such delineation is essential not only to establish the regulatory architecture but also to perform a systems level analysis of the network. The tryptophan system has been quantified through various mathematical models[6,18,24], which have been verified with experimental data. The model equations for the process and the regulators are also depicted in Fig. 2.

**Fig. 2.**
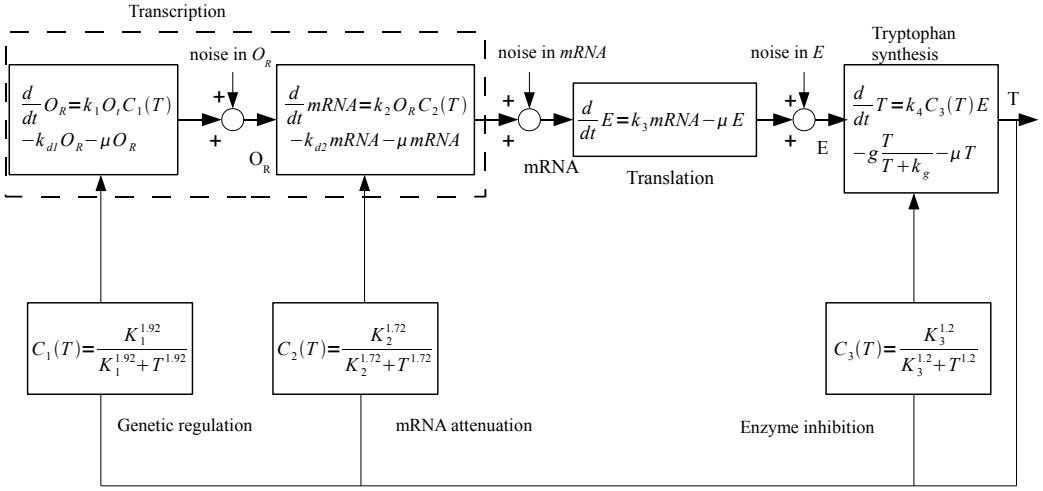
The tryptophan regulatory system in *E. coli* [24]

A question that arises in regards to the design is that, why has the tryptophan system evolved with multiple levels of regulation. It is apparent from Fig. 2 that seven alternate regulatory architectures (one with all three feedback loops, three with dual feedback loops and three with single feedback loops) are possible. From a physiological perspective, a genetic network responds to external cues by regulating the synthesis of protein. This ensures that the system responds by synthesizing the appropriate amounts of protein. Absence of genetic regulation would result in wasteful synthesis of protein thereby imposing additional burden during starvation. Thus, a regulatory design that does not include *C*_1_ is not physiologically relevant. It has been shown by Chaudhary et al.[7] that the role of enzyme inhibition (*C*_3_) is insubstantial under starvation. Thus, the (*C*_1_,*C*_2_) dual feedback design and the (*C*_1_, *C*_2_, *C*_3_) multiple feedback design show identical response. Thus, the physiological designs that are relevant are only (*C*_1_) and (*C*_1_, *C*_2_, *C*_3_), which is nearly same as (*C*_1_, *C*_2_) (see [7]). In this context, a linear analysis has shown that the three levels of regulation, (*C*_1_, *C*_2_, *C*_3_), in a tryptophan system of *E.coli* results in a higher bandwidth relative to a single feedback loop design with *C*_1_ alone[24]. Thus, the multiple feedback loop design is responsible for achieving a rapid synthesis of intracellular tryptophan under starved conditions in about five minutes (see solid line in Fig. 3 (a)), whereas the single feedback loop design attains homoeostatic tryptophan levels only in about 50 minutes (dashed line in Fig. 3 (a)). However, it is well-known that the enhanced bandwidth in a linear system also implies poorer noise-mitigation properties. Specifically, the linearised tryptophan system in Fig. 2 will consist of four poles and two zeros, resulting in a roll-off rate of -40dB/decade, while the linearised single feedback loop consists of four poles only, resulting in a roll-off rate of -80 dB/decade at high frequency. Thus, a frequency response analysis of the linearised tryptophan system suggests that the single feedback loop design can effectively mitigate the high frequency component than the multiple feedback loop design.

**Fig. 3.**
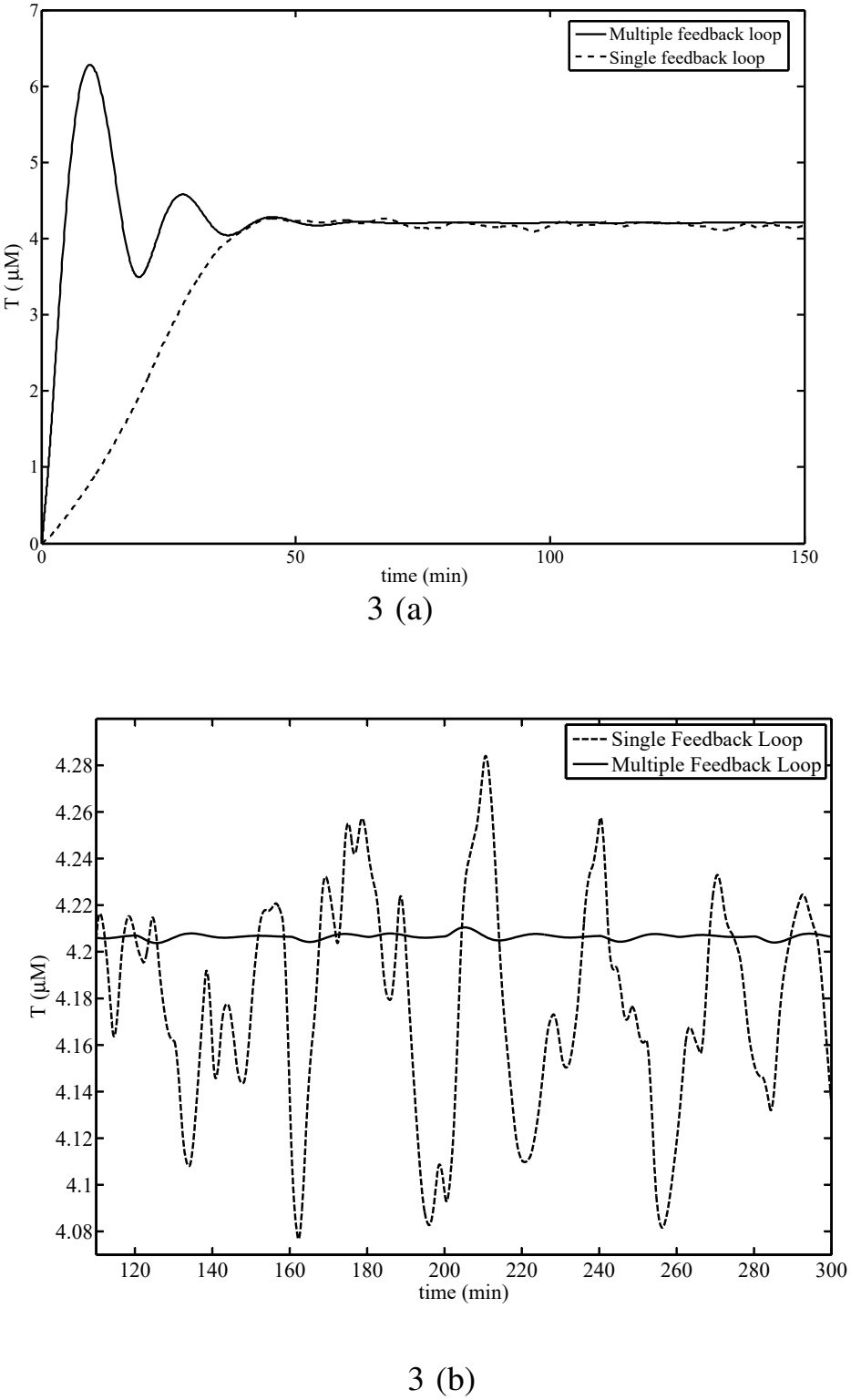
(a) Response of the single and multiple feedback loop strategies to noise in transient state. The solid curve represents the physiological response of the tyrptophan system. The dashed curve represents response of the system without attenuation and inhibition and the genetic regulation parameters altered (*η*_*H*1_ = 0.5 and *K*_1_ = 8 × 10^*−*8^ *µ*M (b) Response of the single and multiple feedback loops when a Gaussian white noise was introduced in mRNA (*σ* = *±* 10%) after both the systems attained steady state.

This work investigates the design elements that limit the adverse impact of multiple nonlinear feedback loop architecture on noise propagation in the tryptophan system. We demonstrate that the noise mitigation properties of the tryptophan system are a direct result of nonlinear regulation. The study suggests that the tryptophan system of *E. coli* has evolved to simultaneously satisfy the two properties of rapid response (high bandwidth) and noise mitigation (high roll off). In particular, the multiple feedback loop is responsible for the rapid response, while the nonlinear regulator, specifically mRNA attenuation, provide effective noise mitigation.

## 3 Results

### 3.1 Performance of tryptophan system in presence of noise

In order to study the dynamic response of the tryptophan system in presence of noise, the model equations shown in Fig. 2 were simulated along with Gaussian white noise injected additively to mimic randomness in genetic regulation, transcription and translation, as indicated in the figure. The performance of the tryptophan system was compared with that of a mutated system consisting of only a single feedback representing genetic regulation. The standard deviation of the noise signal was maintained at 10 % of the steady state concentrations when the bacteria is subjected to complete starvation. Fig. 3 shows the simulated tryptophan concentration. As indicated previously, the multiple feedback loop design synthesises tryptophan rapidly in about 5 minutes, whereas the mutated system takes about 50 minutes to to achieve homoeostatic levels, which is consistent with the higher bandwidth noted for the corresponding linearised system. However, a counter-intuitive picture emerges with regards to noise attenuation. After achieving homoeostatic, the standard deviations of the multiple and single feedback loop systems are found to be 0.001*µ*M and 0.045*µ*M. This result is inconsistent with the linearised multiple feedback loop design of the tryptophan system, where the increased bandwidth corresponds to smaller roll-off rate and have poorer noise mitigation at higher frequency. Hence, the multiple feedback loop design exhibits superior noise attenuation than the mutated single feedback loop system (see Fig. 3 (b)). Thus, it is clear that the nonlinear multiple feedback loop design in the tryptophan system not only exhibits faster response but also robustness to noise. It is therefore natural to question the design principles that endow the tryptophan system with a higher bandwidth as well as a superior noise mitigation property.

### 3.2 Performance of mutated tryptophan system using sub-sensitive regulators

A key difference between the tryptophan system shown in the block diagram of Fig. 2 and a linear feedback system lies in the fact that the three regulators (*C*_*i*_(*T*)) are nonlinear and represented by the Hill equation (Eq. 1),

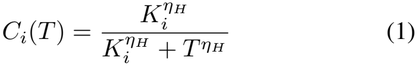

The exponent *η*_*H*_ is termed as the Hill coefficient and *K*_*i*_ is the half saturation constant. The half-saturation constant for the three regulators are *K*_*i*,1_ = 3.53 *µ*M, *K*_*i*,2_ = 0.04 *µ*M and *K*_*i*,3_ = 810 *µ*M. The nature of the nonlinearity is evident from Fig. 4 (a), which depicts the Hill equation for different values of the Hill coefficient on a log scale. The Hill equation is widely used to model feedback in biological regulatory systems and can represent a variety of regulatory behaviour ranging from ultrasensitive to sub-sensitive repression (or activation) depending on the value of the Hill coefficient. A value of zero represents loss of feedback. If *η*_*H*_ *<* 1, the regulator is classified as sub-sensitive, *η*_*H*_ = 1 represents the Michelis-Menten reaction kinetics and *η*_*H*_ *>* 1 signifies that the controller is ultrasensitive. Physiologically, ultrasensitive repression (or activation) is achieved by multiple binding of the repressor (or activator) with a cooperative effect wherein the affinity for binding to subsequent repressor molecules increases multiple times. In order to investigate the role of ultrasensitive regulation in providing effective noise mitigation property to the tryptophan system, we perform the following simulation experiment. The Hill coefficient was modified so that its value for all the three regulators, namely genetic regulation, transcriptional attenuation and enzyme inhibition, was set to 0.5 and the system shown in Fig. 2 was simulated. The tryptophan concentrations corresponding to such sub-sensitive or mildly nonlinear regulation is shown in Fig. 4 (b). The scaled standard deviation for the multiple feedback loop is 2.5 *×* 10^*−*4^, while for the single feedback loop it is 4.3*×* 10^*−*4^. It is clear from the figure, that the multiple feedback loop design using subsensitive regulation yields poorer noise mitigation as compared to a single feedback loop design. This implies that the design principle of combining multiple feedback loop architecture with nonlinear regulation yields a fast response in addition to superior noise mitigation. To assess the role of nonlinear regulation in the multiple feedback loop system, we make use of describing functions to approximate the nonlinear regulators *C*_*i*_(*T*).

**Fig. 4.**
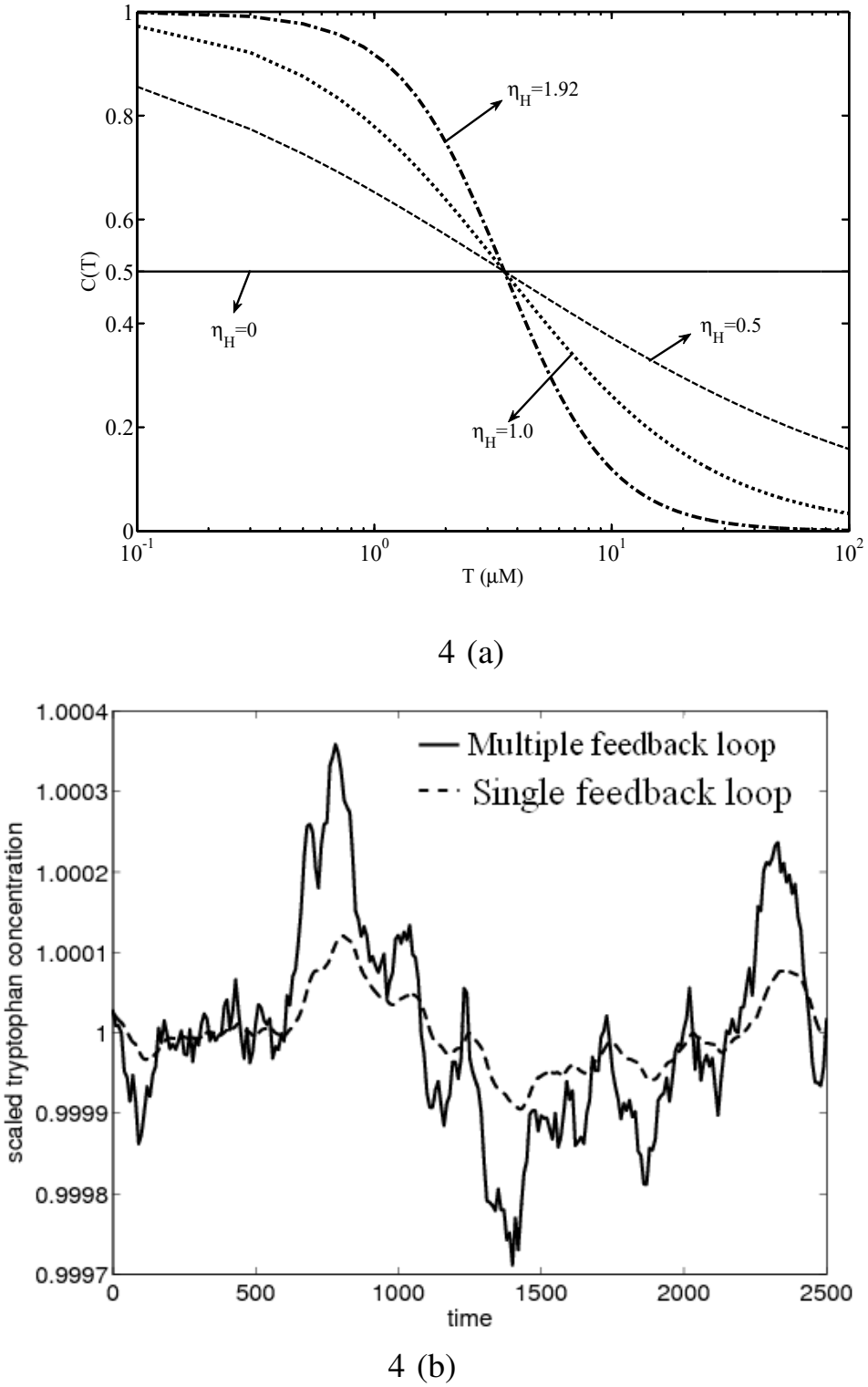
(a) Output characteristics of *C*_1_(*T*) for various values of *η*_*H*_ (b) Comparison of single and multiple feedback loop with sub-sensitive controllers. The multiple feedback loop is worse in noise handling than the single feedback loop

### 3.3 Describing function representation of the nonlinear biological regulators

To maintain homoeostasis, the tryptophan system is tightly regulated in such a manner that when sufficient tryptophan exists in the medium, the three regulators namely genetic regulation, mRNA attenuation and enzyme inhibition repress further synthesis. On the other hand, in absence of tryptophan, the regulators relieve the repressed state of the system enabling rapid tryptophan synthesis. This saturation behaviour is achieved by nonlinear regulators whose variation of gain with intracellular tryptophan for various values of the Hill coefficient is shown in Fig. 4 (a). As seen from the figure, the regulators exhibit significant nonlinearity for a value of Hill coefficient greater than unity even in a narrow range of tryptophan concentration. The Hill equation represents an explicit, static, single-valued smooth nonlinearlity. Fig. 5 (a) shows the plot of the deviation form of the Hill Equation, for all the three regulators, with respect to deviation of tryptophan concentration from its physiological steady-state of 4.2 *µ*M on a linear scale. The negative values of tryptophan concentration represent the lean region, where the corresponding repressive action of the regulator, especially mRNA attenuation, is low, while positive values of the concentration correspond to the rich region where the repressive action by the regulator, especially mRNA attenuation, is strong. From Fig. 5 (a) it can be seen that regulation through mRNA attenuation, *C*_2_(*δT*), exhibits a significant nonlinearity around the homoeostatic level of tryptophan. On the other hand, the extent of inhibition of enzyme at different concentrations of tryptophan, *C*_3_(*δT*), is almost identical, indicating a near absence of the role of feedback. To study the effect of the Hill coefficient in genetic regulation and mRNA attenuation on noise mitigation, we simulated the tryptophan system at different value of *η*_*H*1_ and calculated the mean of the standard deviation obtained using ten different realisations of the Gaussian noise with 1 % of steady state value as standard deviation. In order to ensure that the tryptophan concentration of the thus perturbed system corresponds to the physiological value of 4.2 *µ*M, the half saturation constant *K*_*i*1_ was modified accordingly. The results are shown in a graphical form in Fig. 5 (b). It is evident that as the mRNA attenuation becomes increasingly ultrasensitive, the noise mitigation in the output is more effective. As can be seen from Fig. 5 (b), the regulator *C*_1_(*δT*) plays a marginal role in noise mitigation.

**Fig. 5.**
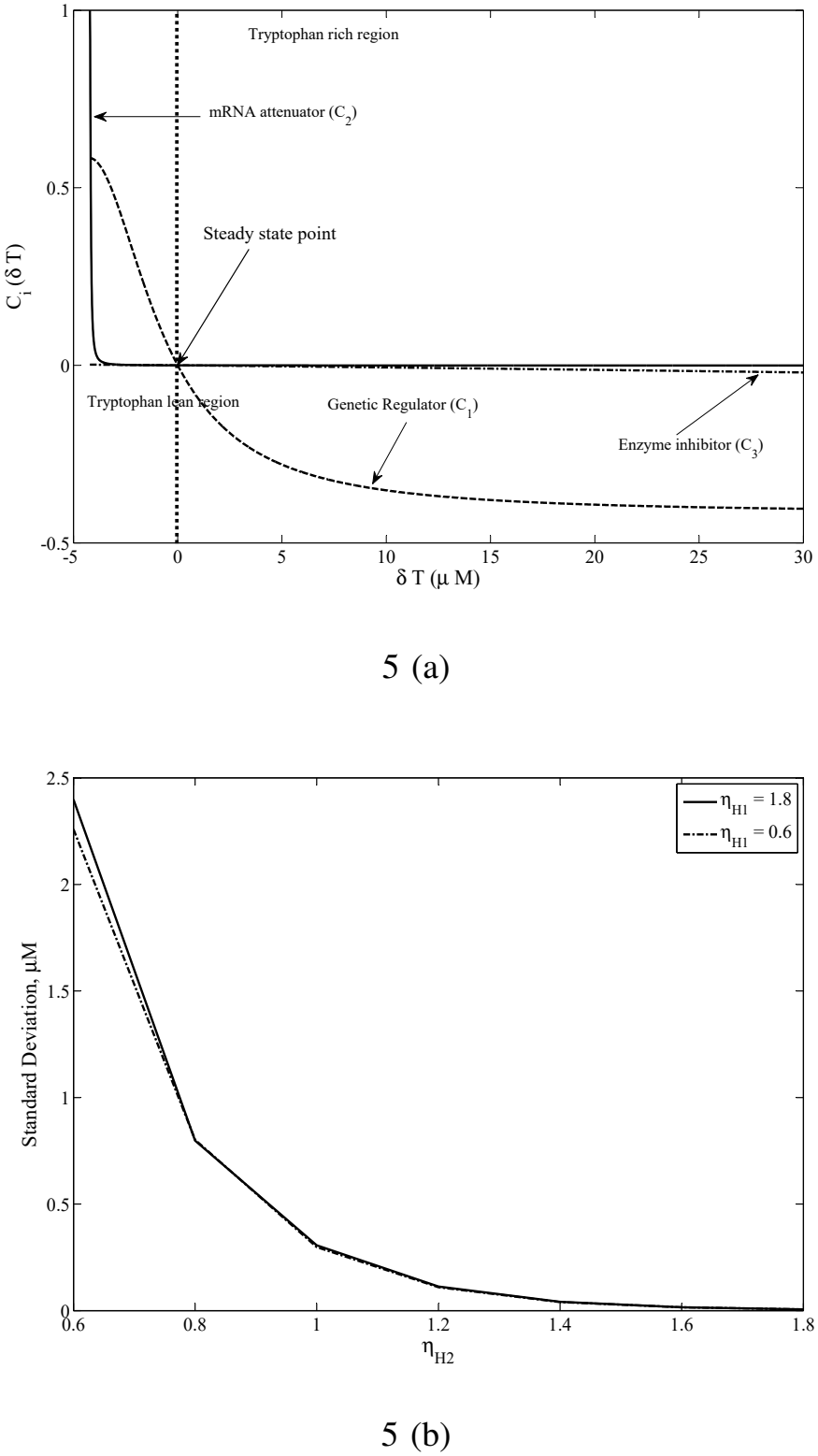
(a) Output of the three regulators in deviation form w.r.t. deviation in tryptophan concentration from homoeostatic level of 4.2 *µ*M.(b) Standard deviation in tryptophan output as a variation of *η*_*H*2_

Frequency response analysis in linear systems is a well-established approach that provides insights on the system bandwidth as well as into the system behaviour in response to high frequency noise. A straightforward extension of the frequency response for nonlinear systems does not exist. However, a heuristic approach that uses quasi-linear input-output models of a nonlinear system called describing functions(DF) enables use of frequency domain methods and intuition for design and analysis of nonlinear systems. In particular, the sinusoidal input describing functions(SIDF) provide a characterisation of the amplitude and frequency dependent effects of the system nonlinearity and can be viewed as an extension of the transfer function concept to nonlinear systems. More importantly, the SIDF provides a better approximation to nonlinearities than the small-gain linearised model based on Taylor’s series expansion. The basic idea in describing functions approach is to replace each nonlinearity in the system with a quasi-linear term whose gain is a function of the amplitude of the sinusoidal input. SIDFs have been widely exploited for controller synthesis for nonlinear systems as well as determination of limit cycles[21]. Here, we use the describing function approach to quasi-linearise the non-linear regulators represented by the Hill equations and use these approximations to obtain a frequency response for the tryptophan system. Since the describing functions technique is based only on the fundamental harmonic of the system, the approximation is valid in closed loop systems only if the system satisfies a filter hypothesis which attenuates the impact of higher harmonics[10]. Since the processes of transcription, translation and synthesis include degradation and dilution of the species due to growth, which act as a low-pass filter (see Fig. 2), the filter hypothesis is satisfied by the tryptophan system and thus describing functions should provide a good representation of the gain characteristics of the three regulators.

We use two ideas to simplify the describing function analysis system multiplicatively. Ideally, the regulator along with the of the nonlinear tryptophan system. Firstly, as seen from Fig. 2 the nonlinear regulator output signals enter the system multiplicatively. Ideally, the regulator along with the process block should constitute the nonlinearity. However, we linearise the bilinear term and focus our analysis only on the nonlinearity in the three regulators. Secondly, the nonlinearity represented by the Hill equation is asymmetric. While there exist methods to obtain describing functions approximation for a general nonlinearity, we choose the simpler approach of accounting for the asymmetric nature of the nonlinearity by dividing the describing function approximation into two parts-a) one that provides a quasi-linear approximation of the control actions when the system is lean in tryptophan (instantaneous concentration less than steady state concentration (see Fig. 6 (a))) and b) the other when the system is rich in tryptophan (see Fig. 6 (b)). Note that the two nonlinearities have been made odd by extending their domains. Methods for obtaining frequency responses of feeedback systems with a static nonlinearity are described in [10] and are based on graphical, analytical and numerical approaches. In this work, we use the graphical method that has been outlined in Appendix C. The block diagram for the tryptophan system as in Fig. 2 with the describing functions *N*_*i*_(*A*) for the three nonlinear regulators (*C*_*i*_(*T*)) is shown in Fig. 7 (a). The processes *G*_*i*_ are the Laplace-domain representation of the four sub-processes of the tryptophan system (see Appendix A for details). The solid lines in Fig. 8 (b) and 8 (a) show the frequency response of the tryptophan system using ultrasensitive regulators as described in Fig. 2, when the system is lean and rich in tryptophan, respectively. These frequency responses are compared to a mutated tryptophan system wherein the three regulators are subsensitive with all three Hill coefficient exponents being set to 0.5.

**Fig. 6.**
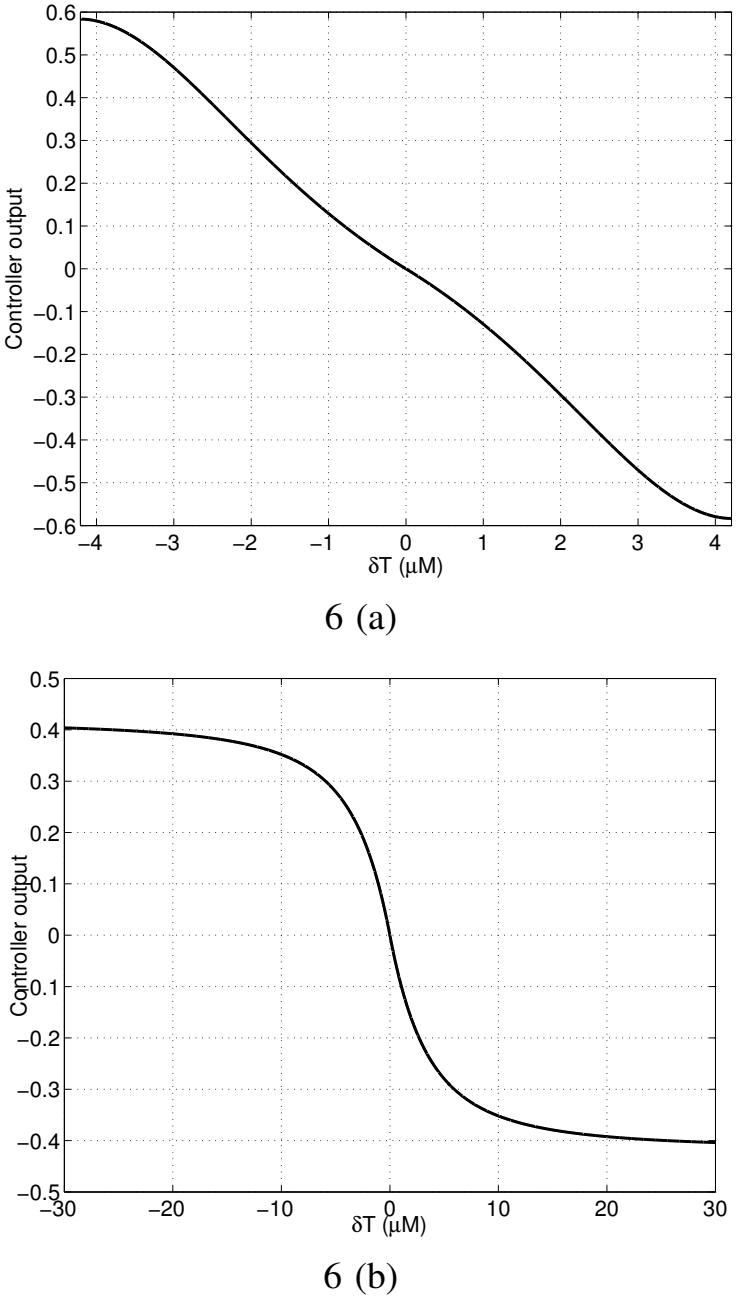
(a) The symmetric depiction of ultrasensitive nature of the nonlinearity in tryptophan lean region (b) Symmetric depiction of the highly repressive nature of the nonlinearity in the tryptophan rich region

**Fig. 7.**
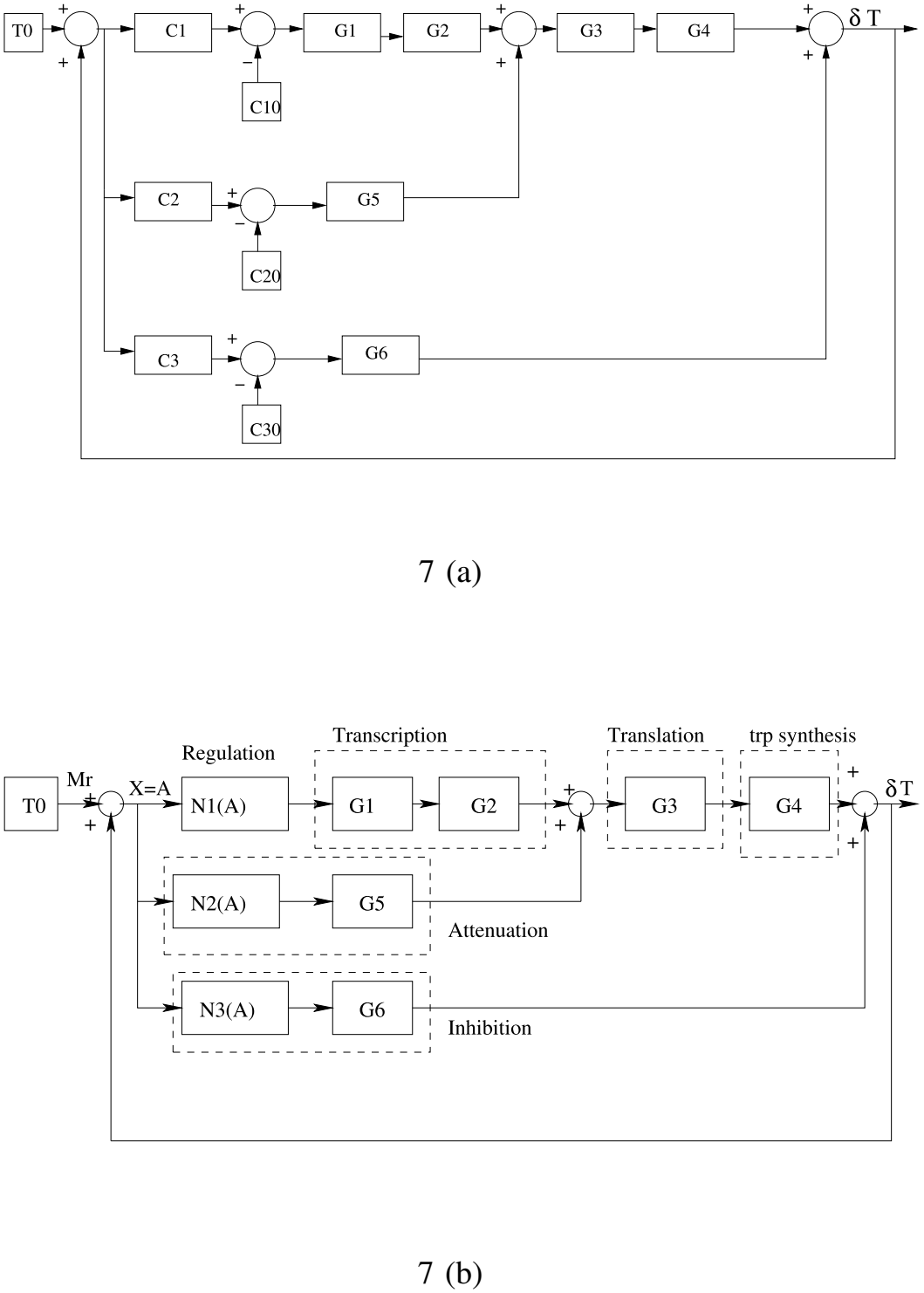
(a) The quasi-linear tryptophan system (b) The SIDF equivalent of tryptophan system. 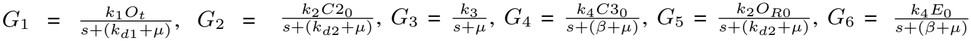

**Fig. 8.**
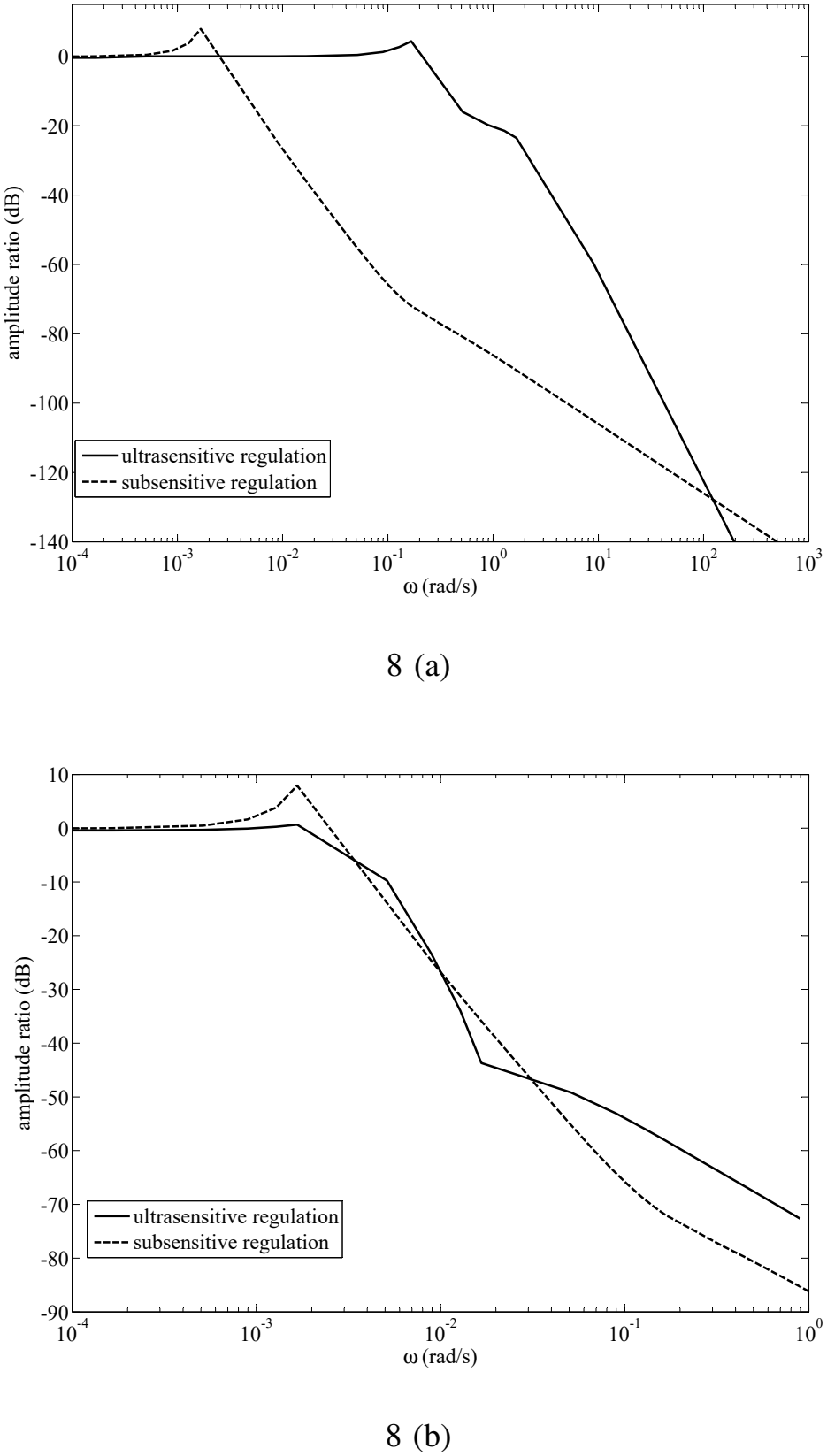
(a) Comparison of frequency response of the tryptophan system with sub-sensitive and ultrasensitive controllers when system is lean in tryptophan (b) Comparison of frequency response of the tryptophan system with sub-sensitive and ultrasensitive controllers when system is rich in tryptophan

When the system is lean in tryptophan (Fig. 8 (a)), with ultrasensitive regulators, the system has a higher bandwidth (about 0.2 rad/s), when compared to the system with subsensitive regulators (about 0.002 rad/s). Thus, the system would transmit signals of frequencies lower than the bandwidth of 0.8 rad/s and attenuate signals of higher frequency. It has been shown in the context of chemotaxis in *E. coli* that the cut-off frequency is 3.4 rad/s for the signalling protein[3]. Thus, the noise component in *E. coli* corresponds to frequencies in a range larger than tens of rad/s. It can be, therefore, expected that the frequency of transcriptional noise will be of the order of magnitudes higher than that in the protein network. The performance of the system when subjected to noise is critical when the system is deficient in tryptophan. The presence of few regulator molecules, in such a scenario, increase the significance of the binding and unbinding of tryptophan to operator, mRNA and enzyme. The system with ultrasensitive controllers has a higher roll-off rate (-61 dB/decade) as compared to the system with sub-sensitive controllers (-22 dB/decade). The higher roll-off rate indicates that the ultrasensitive controllers are more effective in noise mitigation as compared to the sub-sensitive controllers. The amplitude ratios of the subsensitive and ultrasensitive design at the corresponding peak resonance frequency is noted as 7.92 dB and 4.34 dB respectively. When the system is rich in tryptophan, it can be seen from Fig. 8 (b) that the system with sub-sensitive controllers has almost the same bandwidth as the system with ultrasensitive regulators (approximately 0.002 rad/s). However, the system with sub-sensitive controllers shows greater degradation in performance in the higher frequency region than the system with ultrasensitive controllers. This is evident from the higher peak resonance exhibited by the frequency response curve of the system with sub-sensitive controllers. The peak resonance with sub-sensitive regulators is around 8 dB, while with ultrasensitive regulators it is around 0.65 dB. It is well known that a lower resonance peak implies better relative stability of the system.

In summary, the frequency response analysis of the tryptophan regulatory system indicates that the multiple feedback loops are responsible for the small rise time of the system when the system is exposed to starvation stress, while the ultrasensitivity of the regulators is responsible for effective noise attenuation. The rise time and noise inhibition are, both, critical for the tryptophan system. The multiple feedback loop, coupled with ultrasensitive regulators achieve both the objectives– faster response and effective noise mitigation– for the tryptophan system in *E. coli*.

## 4 Discussion

When faced with starvation of tryptophan, the bacterium *Escherichia coli*, synthesises the essential amino acid rapidly in less than five minutes. This rapid synthesis is essential for its survival. On the other hand, when the intracellular tryptophan concentration attains homoeostatic levels, either due to synthesis or uptake from a tryptophan rich medium, the synthesis of tryptophan is repressed. Moreover, the system must maintain homoeostatic levels in presence of uncertainty. The homoeostatic level is determined by evolutionary process and deviation from the optimal level may have adverse effect on growth and survival of the organism. Thus, the system must maintain tryptophan concentration in a narrow band even in a highly uncertain environment. Therefore, the organism needs to be highly sensitive to nutritional requirement, but insensitive to noise. Thus, the design must trade off between fast response and uncertainty. Such a phenotypical response is achieved through a multiple feedback loop design, with the three regulators, used to describe genetic regulation, mRNA attenuation and enzyme inhibition, being ultrasensitive. The overall design principle is a combination of multiple feedback loops and ultrasensitive nonlinear regulation.

The multiple feedback loop design has been shown to increase the bandwidth of the system, allowing to react to high frequency inputs thereby exhibiting rapid synthesis of tryptophan. It is well known that a higher bandwidth in a linear feedback system, results in smaller roll-off rates. Thus, a linear feedback system with a high bandwidth will also react to high frequency noise, thus showing poor noise mitigation. Our work demonstrates that when the system is lean in tryptophan, the ultrasensitive regulation relieves repression, thereby providing higher bandwidth as well as higher roll-off rates relative to a subsensitive regulator design. This rapid synthesis as well as high frequency noise mitigation provides enhances the ability of the organism to survive. The ultrasensitive multiple feedback loop present in the tryptophan regulatory system satisfies the requirements of a good feedback structure– high gain in low frequency region (which is characterised by a higher bandwidth) and a faster roll off rate in the high frequency region (an indicator of noise mitigation). It is important to characterise the noise in a gene circuit in terms of system goals. In context of the tryptophan system, the cut-off frequency beyond which the components are detrimental to the existence of the organism need to be experimentally established. Further, the levels of attenuation at high frequencies needs to be determined.

The tryptophan system used in this work, is prototypical of regulatory networks used in system biology. The Hill equation is commonly used to represent regulation in the genetic, signalling and metabolic pathways. The describing function approach can therefore be gainfully used to analyse biological systems.

## Authors Note

The research presented in this manuscript was carried out during the mid-2000s, primarily as part of an M.Tech research project. This manuscript was created at the end of the research but remained unpublished. We have now decided to make the original manuscript publicly available through bioRxiv to preserve the research and provide access to its mathematical formulation, analysis, and findings as an archival record.

## A A transfer function representation of the tryptophan system

A frequency domain analysis gives valuable information of the system performance in the low and high frequency regions. It can be seen from Fig. 2, the dynamic equations describing the tryptophan synthesis process are of the following nature-

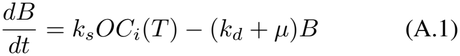

where, *B* is the concentration of species being synthesised (free operator, mRNA, enzyme and tryptophan),

*k*_*s*_, rate of synthesis of *B*,

*O*, concentration of species used for synthesis of *B*,

*k*_*d*_, rate of degradation of *B*,

*µ*, rate of growth of organism and

*C*_*i*_(*T*), nonlinear controller.

Equation A.1 is bilinear in terms of *O* and *C*_*i*_(*T*). We use a Taylor’s series expansion to convert the bilinear term as a series of two terms–

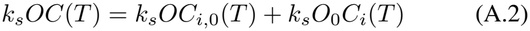

where, *O*_0_ is the value about which the system is quasi-linearised and *C*_*i*,0_(*T*) is the value of *C*_*i*_(*T*) at *T* = *T*_0_.

Using Eq. A.2, Eq. A.1 can now be written as

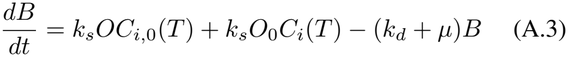

Taking the Laplace transform of Eq. A.3,

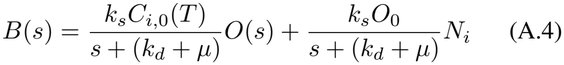

The SIDF can be viewed as the amplitude and phase relationship between the input sinusoid signal and the fundamental harmonic of the output. The term *N*_*i*_ is the describing function representation for the nonlinear term *C*_*i*_(*T*), which represents approximately the transfer characteristics of the nonlinearity for a sinusoidal input and is termed sinusoidal input describing functions (SIDF)[10]. The tryptophan system satisfies the filter hypothesis as the growth and degradation kinetics of *B* act as a low pass filter (for details, see [10]). The following section describes the steps involved in frequency response analysis using describing functions.

### A.1 The SIDF for the the three regulators

The three regulators namely, genetic regulation, mRNA attenuation and enzyme inhibition are represented by the Hill equation, Eq. 1. The salient feature of this nonlinearity is its asymmetric nature as seen in Fig. 5 (a). To account for the asymmetric nature of the nonlinearity, we divide the describing function formulation into two parts– a) when the system is lean in tryptophan (see Fig. 6 (a)) and b) the other when the system is rich in tryptophan (see Fig. 6 (b)).

### A.2 The frequency response of the tryptophan system

To formulate the frequency response problem, the system is quasi-linearised around tryptophan concentration, *T* = 4.2 *µ*M, as discussed in the above section. The SIDF equivalent of the nonlinear system is shown in Fig. 7 (b). Table A.1 lists the details of the parameters in these equations. While deriving the equations for the SIDF of the Hill equations, the output from the final transfer function, *G*_4_ (in Fig. **??**7) is used as a feedback signal to the nonlinear controller, *C*_1_. The reference signal is considered to be zero. Since the Hill equation requires tryptophan concentration in absolute units, the SIDF for the three controllers, namely. genetic regulation (*C*_1_), mRNA attenuation (*C*_2_) and tryptophan inhibition (*C*_3_) was derived by considering the nonlinearity to be

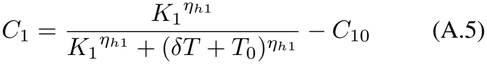

where,

*T*_0_ = tryptophan concentration at steady state = 4.2 *µ*M *δT* = deviation in tryptophan concentration from *T*_0_ *C*_10_ = controller output at *δT* = 0.

The formulation of the SIDF for frequency response analysis is mentioned in detail in Appendix B.

**Table A.1.** Parameters of the transfer function model.

| Parameter | Definition | Value |
|---|---|---|
| *k*<sub>1</sub> | Kinetic rate constant for free operator synthesis | 50 min<sup>−1</sup>, |
| *k*<sub>2</sub> | Kinetic rate constant for mRNA synthesis | 15 min<sup>−1</sup> |
| *k*<sub>3</sub> | Kinetic rate constant for enzyme synthesis | 90 min<sup>−1</sup> |
| *k*<sub>4</sub> | Kinetic rate constant for tryptophan synthesis | 59 min<sup>−1</sup> |
| *μ* | specific growth rate | 0.01 min<sup>−1</sup> |
| *k*<sub>*d*1</sub> | degradation constant of *O*<sub>*R*</sub> | 0.5 min<sup>−1</sup> |
| *k*<sub>*d*2</sub> | degradation constant of mRNA | 15 min<sup>−1</sup> |
| *g* | kinetic constant for tryptophan uptake | 25 *μ*M min<sup>−1</sup> |
| *O*<sub>*t*</sub> | total operator site concentration | 3.32 nM |
| *K*<sub>*g*</sub> | half saturation constant for tryptophan uptake | 0.2 *μ*M |
| *β* | $\frac{gK_g}{(T_0+K_g)^2+\mu}$ | - |

## B SIDF equations for regulation, attenuation and inhibition controllers

For the formulation of the SIDF each controller is approximated as a gain changing nonlinearity. All the nonlinearities encountered in the tryptophan system are static nonlinearities, i.e. their structure is independent of the frequency of input.

### B.1 SIDF for genetic regulation(C_1_)

For the genetic regulation, the SIDF is

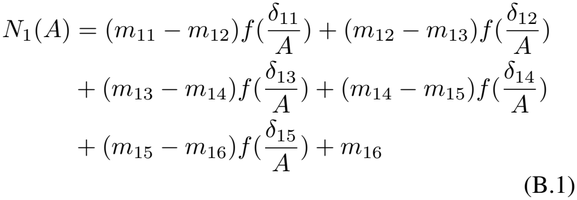

where,

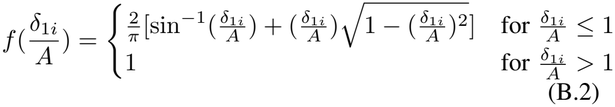

*δ*_1*−i*_ = abscissa break points, i.e. point at which change in slope occurs for *N*_1_, *m*_1*i*_ = slope occurring in the range *δ*_*i* − 1_ *< x≤ δ*_*i*_ for *N*_1_, *A* = amplitude of input to the nonlinearity

### B.2 SIDF for mRNA attenuation(C_2_)

For mRNA attenuation, the SIDF is

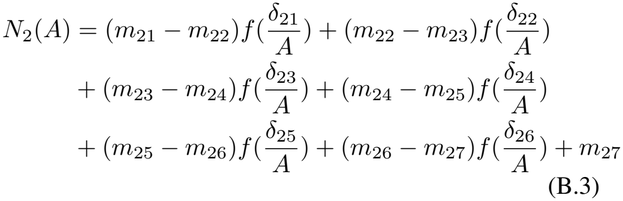

where, 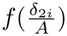 is obtained by Eq. B.2. Rest all symbols have same meaning as for *N*_1_.

### B.3 SIDF for enzyme inhibition(C3)

For enzyme inhibition the SIDF is

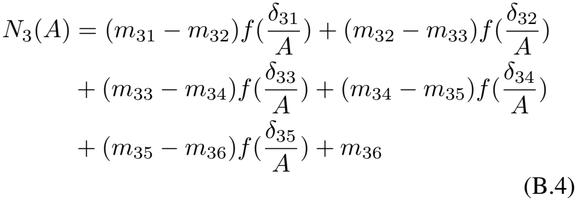

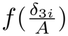 is obtained by Eq. B.2.

## C Derivation of the frequency response equations for the tryptophan regulatory system[10]

The process transfer functions in Figs. 7 (a) and 7 (b) are 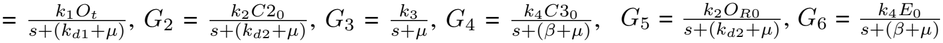.

### C.1 Frequency Response Analysis for the multiple feedback loop

The relation between the output, namely tryptophan concentration *T* and reference input *T*_0_ in Fig. **??**(7b) is given by the following loop transfer function

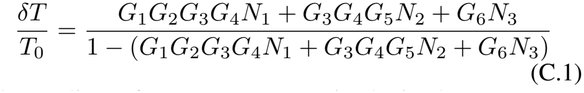

The nonlinear frequency response is obtained as

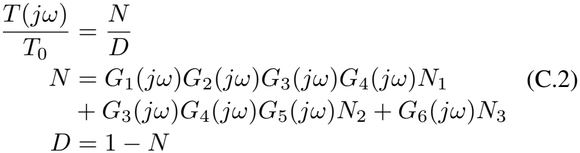

But, the value of the feedback signal *X*(see Fig. 7b) (and therefore the value of *N*_1_, *N*_2_ and *N*_3_) is not immediately obtainable for a given value of *T*_0_. It is clear from Fig. 7b, that the transfer function relating *X* and *T*_0_ is given as

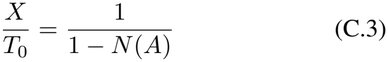

Since the RHS is also dependent on *X*, the above equation can be solved graphically.

#### C.1.1 Derivation of the frequency response equations

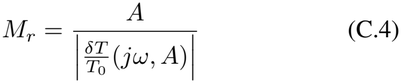

where *M*_*r*_ is the amplitude of the reference signal *T*_0_.

To enable formulation for graphical calculation let, 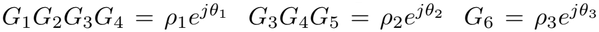 and *N*_1_ = *ρ*_*N*1_ *N*_2_ = *ρ*_*N*2_ *N*_3_ = *ρ*_*N*3_

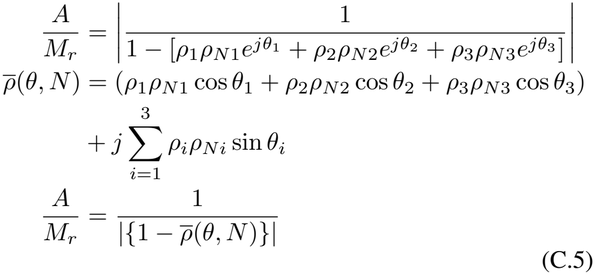

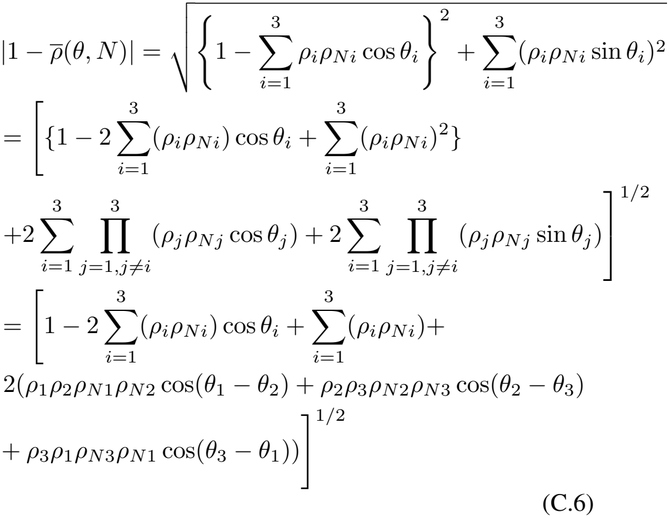

Re-arranging Eq. C.5,

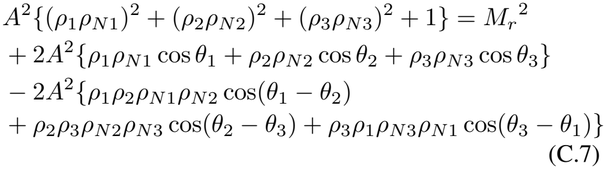

Solving the above equation, we get

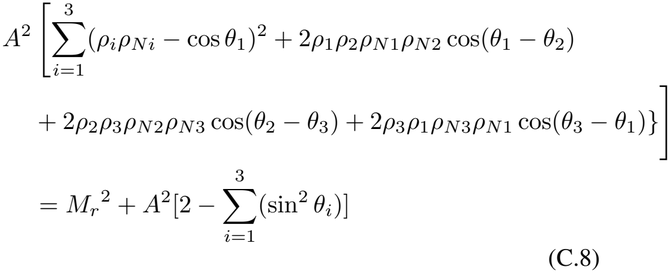

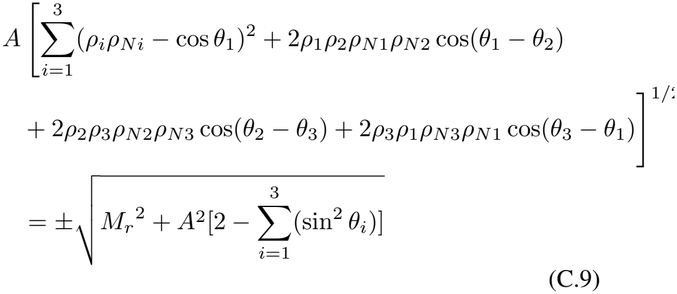

### C.2 Graphical solution for frequency response

From equation C.9 it is clear that the left hand side depends on the describing functions, while the right hand side does not. The describing functions *N*_1_, *N*_2_ and *N*_3_ are functions of frequency(*ω*) and amplitude(*A*). On the other hand, the RHS of Eq. C.9 depends only on the amplitude (*A*) of the SIDF. Thus, there are two unknowns (*ω* and *A*) but only one equation (Eq. C.9) to solve the frequency response problem.

For a fixed value of *ω*, we can choose various values of *A*. The left hand side of Eq. C.9 is plotted against various values of *A*. Similarly, the right hand side of Eq. C.9 is also plotted for the same set of values of *A*. Fig. C.1 shows such a plot for *ω* = 0.00167 rad/s and *Aϵ*[*− T*_0_, *T*_0_]. The points of intersection of the two curves give the value of *A* for a particular *ω*. This value can then be substituted in Eq. C.2 to get the amplitude ratio for a particular *ω*.

**Fig. C.1.**
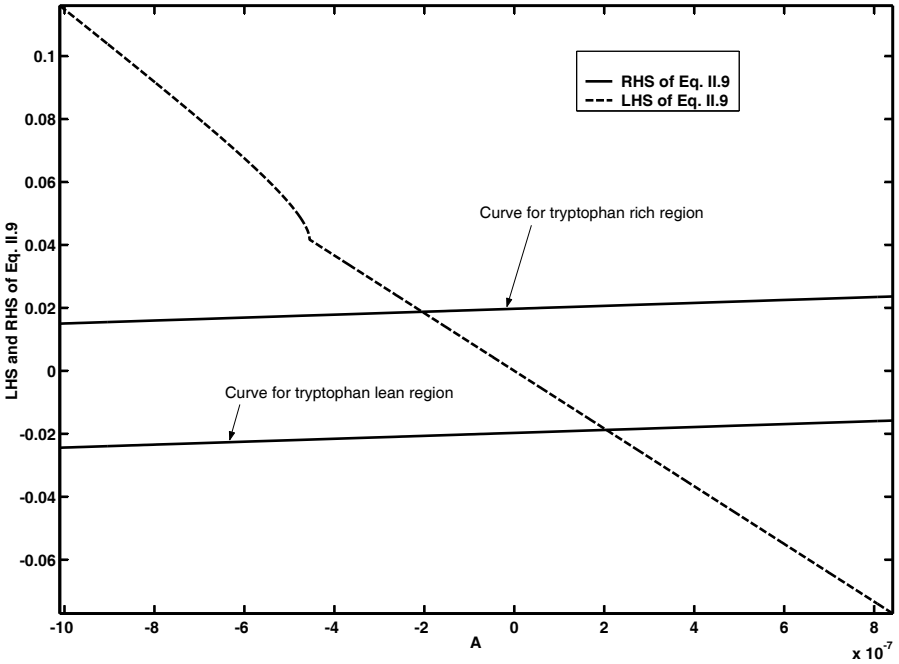
Graph of LHS and RHS of Eq. C.9 for *ω* = 0.00167 rad/s

